# Intratumoral Heterogeneity of Tumor Infiltration of Glioblastoma Revealed by Joint Histogram Analysis of Diffusion Tensor Imaging

**DOI:** 10.1101/187450

**Authors:** Chao Li, Shuo Wang, Jiun-Lin Yan, Rory J. Piper, Hongxiang Liu, Turid Torheim, Hyunjin Kim, Jinjing Zou, Natalie R. Boonzaier, Rohitashwa Sinha, Tomasz Matys, Florian Markowetz, Stephen J. Price

**Affiliations:** Cambridge Brain Tumor Imaging Laboratory, Division of Neurosurgery, Department of Clinical Neurosciences, University of Cambridge, Addenbrooke’s Hospital, Cambridge, UK (CL, JLY, RJP, NRB, RS & SJP); Department of Neurosurgery, Shanghai General Hospital, Shanghai Jiao Tong University School of Medicine (originally named “Shanghai First People’s Hospital”), China (CL); Department of Radiology, University of Cambridge, Cambridge, UK (SW & TM); Department of Neurosurgery, Chang Gung Memorial Hospital, Keelung, Taiwan (JLY); Chang Gung University College of Medicine, Taoyuan, Taiwan (JLY); Molecular Malignancy Laboratory, Hematology and Oncology Diagnostic Service, Addenbrooke’s Hospital, Cambridge, UK (HL); Cancer Research UK Cambridge Institute, University of Cambridge, Cambridge, UK (TT, HK & FM); CRUK & EPSRC Cancer Imaging Centre in Cambridge and Manchester, Cambridge, UK (TT & FM); Statistical laboratory, Centre for Mathematical Sciences, University of Cambridge, UK (JZ); Developmental Imaging and Biophysics Section, Institute of Child Health, University College London, London, UK (NRB); Cancer Trials Unit Department of Oncology, Addenbrooke’s Hospital, Cambridge, UK (TM); Wolfson Brain Imaging Centre, Department of Clinical Neurosciences, University of Cambridge, Cambridge, UK (SJP)

**Keywords:** Glioblastoma, Magnetic Resonance Imaging, Diffusion Tensor Imaging, Heterogeneity

## Abstract

**Introduction:** Glioblastoma exhibits profound tumor heterogeneity, which causes inconsistent treatment response. The aim of this study was to propose an interpretation method of diffusion tensor imaging (DTI) using joint histogram analysis of DTI-p and-q. With this method we explored the patterns of tumor infiltration which causes disruption of brain microstructure, and examined the prognostic value of tumor infiltrative patterns for patient survival.

**Materials and methods:** A total of 115 primary glioblastoma patients (mean age 59.3 years, 87 males) were prospectively recruited from July 2010 to August 2015. Patients underwent preoperative MRI scans and maximal safe resection. DTI was processed and decomposed into p and q components. The univariate and joint histograms of DTI-p and-q were constructed using the voxels of contrast-enhancing and non-enhancing regions respectively. Eight joint histogram features were obtained and correlated with tumor progression and patient survival. Their prognostic values were compared with clinical factors using receiver operating characteristic curves.

**Results:** The subregion of increased DTI-p and decreased DTI-q accounted for the largest proportion. Additional diffusion patterns can be identified via joint histogram analysis. Particularly, higher proportion of decreased DTI-p and increased DTI-q in non-enhancing region contributed to worse progression-free survival (hazard ratio = 1.08, *p*< 0.001) and overall survival (hazard ratio = 1.11, p < 0.001).

**Conclusions:** Joint histogram analysis of DTI can provide a comprehensive measure of tumor infiltration and microstructure change, which showed prognostic values. The subregion of decreased DTI-p and increased DTI-q in non-enhancing regions may indicate a more invasive habitat.

## Introduction

Glioblastoma is the commonest primary malignant tumor in the central nervous system of adults(Ricard et al., 2012). It is among the most lethal cancers, characterized by diffuse infiltration into the normal brain tissue (Weller et al., 2017), which renders a total resection impossible. Progression after surgery is almost inevitable, and predominantly manifest adjacent to the resection cavity(Giese et al., 2003).

It is recognized that glioblastoma is heterogeneous in its infiltrative pattern. Some tumors may disseminate into the surrounding brain tissue and change the brain microstructure earlier and more intensively (Price et al., 2007). Recent genomic findings have revealed that multiple tumor clones co-exist in the same tumor and may display diverse biological traits (Sottoriva et al., 2013; Verhaak et al., 2010). Migratory clones may contribute to a more diffuse and invasive phenotype, which is thought to be especially responsible for treatment failure (Giese et al., 2003). Therefore, understanding the intratumoral heterogeneity of glioblastoma infiltration is of clinical significance for targeted surgery and radiotherapy.

Magnetic resonance imaging (MRI) has unique advantages in understanding spatial variations within glioblastoma. Current clinical management is primarily based on structural sequences, among which the contrast enhancement on post-contrast T1-weighted imaging is most widely-used. However, these findings are limited since it just reflects the failure of blood brain barrier. Other sequences, such as fluid attenuated inversion recovery (FLAIR), although have been combined in patient assessment (Wen et al., 2010), but still proved to be non-specific in detecting tumor infiltration (Price et al., 2006).

Diffusion tensor imaging (DTI), a method that measures the magnitude and direction of water molecule movement, has shown its sensitivity in detecting tumor infiltration and microstructure change (Price et al., 2017). Glioblastoma is recognized to preferentially migrate along the white matter tracts and causes disruption to the microstructure of the tracts (Hambardzumyan and Bergers, 2015).The diffusion of water molecules in the tumor core and peritumoral brain tissue is consequently altered. By decomposing the tensor into isotropic (DTI-p) and anisotropic (DTI-q) components, the directional diffusion of water molecules can be measured (Pena et al., 2006). This approach has been identified to be useful in predicting tumor progression (Price et al., 2007) and patient survival (Mohsen et al., 2013). However, it remains to be discovered if integrating these decomposed components can offer a more comprehensive measure of tumor infiltration. Furthermore, several molecular biomarkers, such as isocitrate dehydrogenase (IDH) mutations (Parsons et al., 2008) and oxygen 6–methylguanine-DNA methyltransferase (MGMT) promoter methylation (Hegi et al., 2005) have been reported to be of diagnostic and prognostic significance for glioblastoma. Whether these DTI markers, particularly DTI-p and-q, can provide additional value to molecular markers is still unclear.

The purpose of this study is to propose a novel interpretation method of DTI by using the joint histogram analysis of DTI-p and-q. With this method we explored the heterogeneity of tumor infiltration and examined the prognostic value of tumor infiltrative patterns for patient survival. Our hypothesis was that the joint histogram analysis of DTI-p and-q could provide: 1) a more comprehensive measure of heterogeneity of tumor infiltration and microstructure change; 2) a more accurate detection of infiltrative subregions which are more responsible for tumor progression; and 3) incremental prognostic values for glioblastoma patients when integrated with IDH-1 mutation, MGMT methylation status and other clinical factors.

## Materials and methods

### Patient population

This study was approved by the local institutional review board. Signed informed consent was obtained from all patients. A total of 136 patients with supratentorial primary glioblastoma were prospectively recruited for surgical resection from July 2010 to August 2015. All patients displayed good performance status (World Health Organization performance status 0-1). Exclusion criteria included history of a previous brain tumor, cranial surgery, radiotherapy/chemotherapy, or contraindication for MRI scanning. To achieve maximal safe resection, tumor resection was performed with the guidance of neuronavigation (StealthStation, Medtronic) and 5-aminolevulinic acid fluorescence (5-ALA). After surgery, 115 (84.6%) histologically confirmed glioblastoma patients (mean age 59.3 years, range 22-76 years, 87 males) were included. Twenty-one patients were excluded due to the non-glioblastoma pathology diagnosis.

Chemoradiotherapy was performed when patients were stable after surgery, Extent of resection was assessed according to the postoperative MRI scans within 72 hours, classified as either gross total resection, subtotal resection or biopsy of the contrast enhanced tumor region. Patients were followed up in our neuro-oncology clinics. Treatment response and tumor progression were evaluated according to the Response Assessment in Neuro-oncology criteria (Wen et al., 2010). All MRI and histological data were collected prospectively, whereas survival data were analyzed retrospectively to avoid the issue of pseudoprogression after patients were followed up for > 3 years.

### Pre-operative MRI acquisition

A 3-Tesla MRI system (Magnetron Trio; Siemens Healthcare, Erlangen, Germany) with a standard 12-channel receive-head coil was used for MRI acquisition. MRI sequences included diffusion tensor and anatomical imaging. Diffusion tensor imaging was acquired with a single-shot echo-planar sequence (TR/TE 8300/98 ms; flip angle 90°; FOV 192 × 192 mm; 63 slices; no slice gap; voxel size 2.0 × 2.0 × 2.0 mm; 12 directions; b values: 350, 650, 1000, 1300, and 1600 sec/mm^2^; imaging time: 9 minutes 26 seconds). The anatomical sequences were acquired as following: post-contrast T1-weighted imaging (TR/TE/TI 2300/2.98/900 ms; flip angle 9°; FOV 256 × 240 mm; 176-208 slices; no slice gap; voxel size 1.0 × 1.0 × 1.0 mm,) after intravenous injection of 9 mL gadobutrol (Gadovist,1.0 mmol/mL; Bayer, Leverkusen, Germany); T2-weighted fluid attenuated inversion recovery (FLAIR) (TR/TE/TI 7840-8420/95/2500 ms; refocusing pulse flip angle 150°; FOV 250 × 200 mm; 27 slices; 1 mm slice gap; voxel size of 0.78 × 0.78 × 4.0 mm).

### Imaging processing

For each subject, DTI images were firstly processed with the diffusion toolbox (FDT) (Behrens et al., 2003) of FSL v5.0.8 (FMRIB Software Library, Centre for Functional MRI of the Brain, Oxford, UK) (Smith et al., 2004), during which normalization and eddy current correction were performed. For each voxel of the processed images, the isotropic component (p) and the anisotropic component (q) of DTI were calculated by using the previously described equation (Pena et al., 2006). Anatomical MRI images were coregistered to the processed diffusion tensor images with an affine transformation, using the FSL linear image registration tool (FLIRT) (Jenkinson et al., 2002).

Tumor regions of interest (ROIs) were manually drawn on the post-contrast T1 and FLAIR images using an open-source software 3D slicer v4.6.2 (Fedorov et al., 2012) by a neurosurgeon with > 8 years of experience (CL), and a researcher with > 4 years of brain tumor image analysis experience (NRB) and reviewed by a neuroradiologist with > 8 years of experience (TM). Non-enhancing ROI, defined as the non-enhancing region outside of contrast-enhanced region, were obtained in MATLAB (MathWorks, Inc., Natick MA) by Boolean operations on contrast-enhancing and FLAIR tumor ROIs. For each individual subject, normal-appearing white matter was drawn manually in the contralateral white matter as normal controls. Volumetric analysis of all ROIs was performed in FSL by using the function of fslmaths (Smith et al., 2004). Inter-rater reliability testing was performed using Dice similarity coefficient scores.

### Histogram analysis

Both univariate and joint histogram analysis were performed in the Statistics and Machine Learning Toolbox of MATLAB. The contrast-enhancing and non-enhancing ROIs were analyzed independently. A demonstration of histogram analysis was summarized in Figure 1. Firstly, DTI-p and-q values were obtained from the ROI on a voxel-by-voxel basis. Each tumor voxel value was normalized by dividing it by the mean voxel value of the contralateral normal-appearing white matter. Next, the univariate histograms of DTI-p and-q were constructed from the normalized voxel values using 100 bins within the range 0~5 (Figure 1. A & B). The height of the bins represented the relative frequency of the voxels falling into a specific DTI-p or-q value range. The mean, median, 25th and 75th percentiles of the univariate histogram were calculated.

**Figure 1.**
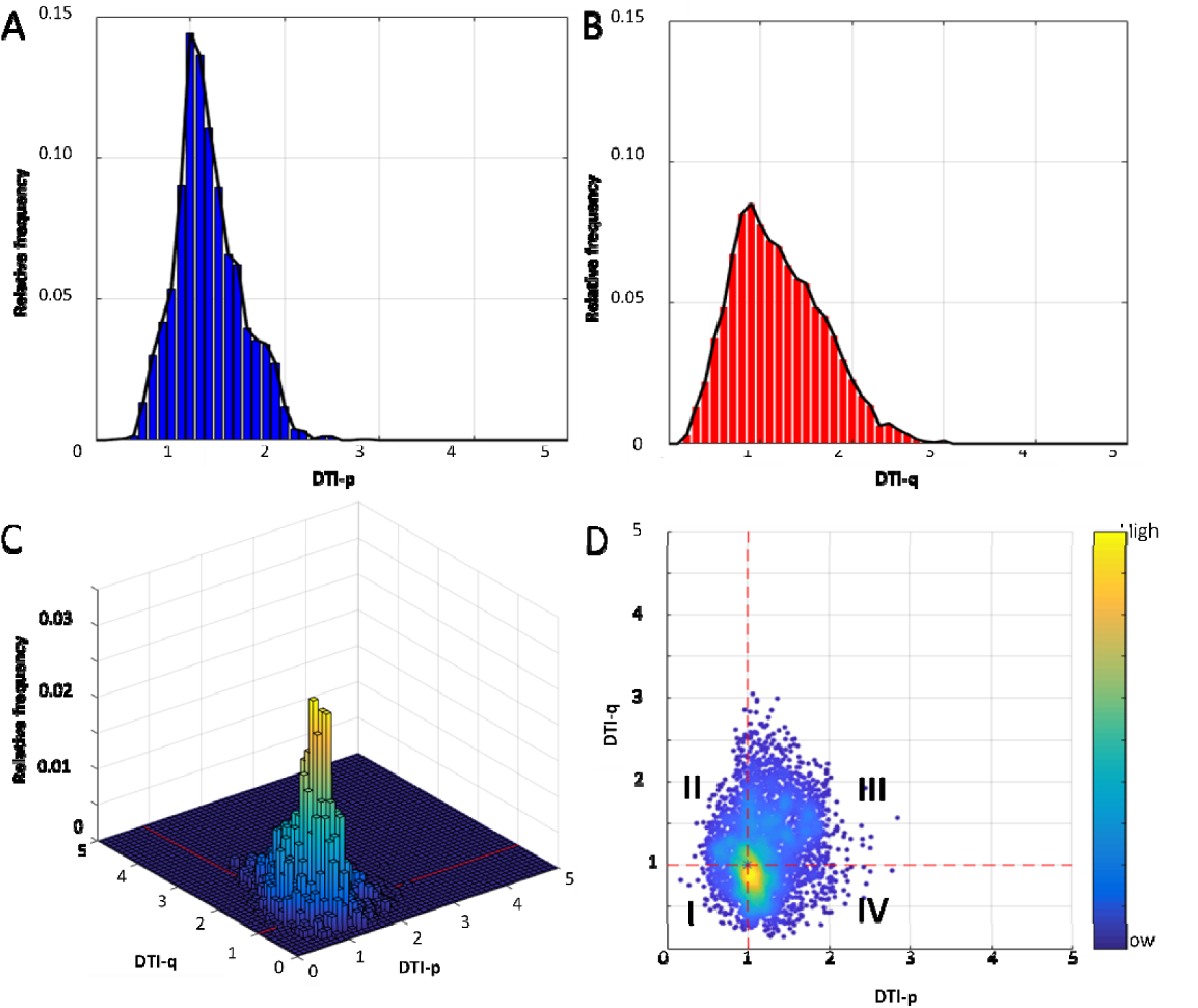
Illustration of the univariate and joint histogram analysis. Univariate histograms of DTI-p (A) and-q (B) were generated using 100 bins. The joint histogram was generated with x-and y-axis representing DTI-p and-q values using 50 × 50 bins. The bin height of the joint histogram represented the relative frequency of voxels in the ROI falling into a specific DTI-p and-q range (C). Four voxel groups of DTI-p and-q abnormalities were obtained (D): I. Voxel Group I (decreased DTI-p/decreased DTI-q); II. Voxel Group II (decreased DTI-p/increased DTI-q); III. Voxel Group III (increased DTI-p/increased DTI-q); IV. Voxel Group IV (increased DTI-p/decreased DTI-q).

The joint histogram was constructed with the x-and y-axis representing the normalized DTI-p and-q values respectively, using 50×50 bins within the range of 0~5 on both axes (Fig 1. C). All voxels from the ROI were assigned to the corresponding histogram bins in the 3D space, according to the DTI-p and-q values they carried. The bin height of the joint histogram represented the relative frequency of the voxels simultaneously falling into a specific DTI-p and-q value ranges. Since the DTI-p and-q values of each voxel have been normalized as above mentioned, the coordinator point (p=1, q=1) was designated to represent the mean diffusion pattern in the contralateral normal-appearing white matter. Thus, according to the relative position to the coordinator point (p=1, q=1), four voxel groups describing the co-occurrence distribution of DTI-p and-q abnormality were obtained (Fig 1. D), namely:

I. Voxel Group I (decreased DTI-p/decreased DTI-q, p↓/q↓)
II. Voxel Group II (decreased DTI-p/increased DTI-q, p↓/q↑)
III. Voxel Group III (increased DTI-p/increased DTI-q, p↑/q↑)
IV. Voxel Group IV (increased DTI-p/decreased DTI-q, p↑/q↓)

The proportion of each voxel group in the ROI (calculated as the summed voxels divided by the total ROI volume) were used as the joint histogram features. These joint histogram features were obtained from both contrast-enhancing and non-enhancing tumor regions, leading to 8 features of each patient.

### Assessment of IDH-1 R132H Mutation and MGMT Methylation Status

IDH-1 R132H mutation was firstly evaluated via immunohistochemistry. After the paraffin was removed and a heat-induced antigen retrieval was performed, IDH-1 R132H mutation-specific antibody (Dianova, Hamburg, Germany) was applied at a 1:20 dilution on slices. A secondary antibody avidin-based detection system was used. In five patients for whom IDH-1 R132H mutation was not detected by immunohistochemistry, tumor DNA was extracted from tumor-rich tissue and sequenced for other rare IDH mutation in codon 132 of the IDH-1 gene and codon 172 of the IDH2 gene using the targeted next generation sequencing (Ion AmpliSeq Cancer Hotspot Panel v2 and (Ion PGM System; Thermo Fisher Scientific).

MGMT promoter methylation status was evaluated as follows: DNA was extracted from the dissected neoplastic cell-rich tissue and was bisulphite-converted using the EpiTect Bisulphite Kit (Qiagen). Pyrosequencing of four CpG sites (CpGs 76-79) in exon 1 of the MGMT gene was then performed by using the CE-Marked therascreen MGMT Pyro Kit on a Pyromark Q24 System (Qiagen). A cut-off of 10% mean methylation for the four CpG sites was used to determine tumors as either methylated or unmethylated based on the published data (Collins et al., 2014; Dunn et al., 2009).

### Evaluation of tumor progression

Tumor progression was diagnosed in our neuro-oncology clinics by the multidisciplinary team. The time to progression was defined as the time period between the surgery date and the date of the first MRI T1W image with contrast that showed tumour progression (as determined by a consultant neuroradiologist). Available tumor progression images were collected and reviewed by three authors (CL, JLY and RJP).

A two-stage semiautomatic coregistration between the progression images and pre-operative postcontrast T1-weighted images was performed using a previously reported tool (van der Hoorn et al., 2016). This coregistration method firstly calculated the transformation matrix between the pre-operative lesion and tumor cavity on progression images. The matrix was then applied to the brain parenchyma (Yan et al., 2017). After coregistration, the progression tumor volume was calculated using FSL function of fslmaths. The progression rate was calculated as the progression tumor volume divided by time to progression.

### Statistical analysis

All analyses were performed in RStudio v3.2.3. Histogram features or tumor volume were tested in Wilcoxon signed rank test due to their non-normal distributions. Cox proportional hazards regression method was performed to evaluate patient survival considering all the other relevant covariates, including IDH-1 mutation status, MGMT methylation status, sex, age, extent of resection, contrast-enhancing tumor volume, together with each univariate and joint histogram feature. Patients who were alive at the last known follow-up were censored. For the Kaplan-Meier analysis, the continuous variables were dichotomized using optimal cutoff values, which were calculated by the R Package “survminer”. Logistic regression models were used to test prognostic values of covariates for 12-, and 18-month overall survival (OS) and progression-free survival (PFS). The baseline models were constructed using all the clinical covariates of IDH-1 mutation status, MGMT methylation status, sex, age, extent of resection, contrast-enhancing tumor volume. Then specific histogram feature was added one by one into the baseline model to assess their incremental prognostic value. The area under the receiver operator characteristics curve (AUC) were compared using one-way ANOVA. Multivariate Cox regression with forward and backward stepwise procedures was performed for the selection of prognostic variables from the clinical factors and the joint histogram features. The forward procedure initiated from the model which only contained one covariate. The backward procedure initiated from the model in which all covariates were included. For each step, the model obtained after adding or deleting one covariate was evaluated using the Akaike Information Criterion (AIC). The final multivariate Cox regression model was constructed using only the features selected by the stepwise procedures. The hypothesis of no effect was rejected at a two-sided level of 0.05.

## Results

### Patients and regions of interest

Among the included 115 patients, 84 (73.0 %) patents received standard dose of radiotherapy plus temozolomide concomitant and adjuvant chemotherapy post-operatively. Other patients, due to their poor post-operative performance status, received short-course radiotherapy (17.4%, 20/115) or best supportive care (9.6%, 11/115). Survival data were available for 80 of 84 (95.2%) patients and 4 (4.8%) patients were lost to follow up. IDH-1 mutation status was available for all patients and 7 of 115 (6.1%) patients were IDH-1 mutant. In the remaining 108 patients, an IDH-1 R132H or other rare IDH-1 and IDH-2 mutations where screened were not detected. MGMT-methylation status was available for 111 patients, among which 48 of 111 (43.2%) patients were methylated. The median OS was 424 days (range 52-1376 days) and the median PFS was 262 days (range 25-1130 days). Patient clinical characteristics are summarized in Table 1. Inter-rater reliability testing of regions of interest (ROIs) showed excellent agreement between the two raters, with Dice scores (mean ± standard deviation [SD]) of 0.85 ± 0.10 and 0.86 ± 0.10 for contrast-enhancing and non-enhancing ROIs respectively. The volumes (mean ± SD) of contrast-enhancing and non-enhancing ROIs were 53.6 ± 33.8 cm^3^ and 62.5 ± 44.0 cm^3^ respectively.

**Table 1.** Clinical characteristics

| **Table 1. Clinical characteristics** |  |
| --- | --- |
| **Variable** | **Patient Number** |
| **Age at diagnosis** |  |
| <60 | 40 |
| ≥60 | 75 |
| **Sex** |  |
| Male | 87 |
| Female | 28 |
| **Extent of resection (of enhancing tumor)** |  |
| Complete resection | 77 |
| Partial resection | 32 |
| Biopsy | 6 |
| **MGMT-methylation status\*** |  |
| Methylated | 29 |
| Unmethylated | 46 |
| **IDH-1 mutation status** |  |
| Mutant | 7 |
| Wild-type | 107 |
| **Tumor volumes(cm3) <sup>#</sup>** |  |
| Contrast-enhancing | 53.6 ± 34.0 |
| Non-enhancing | 62.5 ± 44.2 |
| **Survival (days)** |  |
| Median OS (range) | 424 (52 -1376) |
| Median PFS (range) | 262 (25-1130) |
| *MGMT-methylation status unavailable for 4 patients; <sup>#</sup>mean ± SD of original data. SD: standard deviation; MGMT: O-6-methylguanine-DNA methyltransferase; IDH-1: Isocitrate dehydrogenase 1; cm: centimeters; OS: overall survival; PFS: progression-free survival. |  |

### Diffusion signatures of contrast-enhancing and non-enhancing regions

The diffusion signatures of contrast-enhancing and non-enhancing regions are demonstrated in Table 2. The comparisons of all histogram features between contrast-enhancing and non-enhancing regions showed significance, suggesting the two regions had distinct diffusion patterns. For the univariate histogram features, both contrast-enhancing and non-enhancing regions displayed increased DTI-p (with a value of all greater than 1), compared to normal-appearing white matter, although the increase in contrast-enhancing region is more significant. A consistently decreased DTI-q (with a value of less than 1) was observed in contrast-enhancing region. In contrast, non-enhancing region displayed decreased 25th percentile and median but increased mean and 75th percentile in DTI-q, indicating the non-enhancing region had a heavier tail in DTI-q. Generally, contrast-enhancing region displayed a more significantly increased DTI-p and decreased DTI-q than non-enhancing region.

**Table 2.** Histogram measures

| Table 2. Histogram measures |  |  |  |
| --- | --- | --- | --- |
| Variable | contrast-enhanced region | non-enhancing region | *p* value |
|  | Mean ± SD | Mean ± SD |  |
| DTI-p histogram features |  |  |  |
| 25th percentile | 1.18 ± 0.23 | 1.11 ± 0.15 | < **0.001** |
| Median | 1.47 ± 0.38 | 1.32 ± 0.23 | **0.001** |
| Mean | 1.57 ± 0.36 | 1.37 ± 0.22 | < **0.001** |
| 75th percentile | 1.90 ± 0.60 | 1.59 ± 0.30 | < **0.001** |
| DTI-q histogram features |  |  |  |
| 25th percentile | 0.42 ± 0.14 | 0.71± 0.18 | < **0.001** |
| Median | 0.65 ± 0.19 | 0.95 ± 0.23 | < **0.001** |
| Mean | 0.71 ± 0.19 | 1.01 ± 0.24 | < **0.001** |
| 75th percentile | 0.93 ± 0.24 | 1.24 ± 0.29 | < **0.001** |
| DTI Joint histogram features (%) |  |  |  |
| Voxel Group I | 8.50 ± 10.37 | 5.49 ± 6.17 | < **0.001** |
| Voxel Group II | 3.83 ± 4.92 | 7.27 ± 8.07 | < **0.001** |
| Voxel Group III | 20.78 ± 13.65 | 40.33 ± 18.97 | < **0.001** |
| Voxel Group IV | 66.90 ± 16.28 | 46.92 ± 20.40 | < **0.001** |
| DTI: diffusion tensor imaging; SD: standard deviation; Voxel Group I: decreased DTI-p, decreased DTI-q (p↓/q↓); Voxel Group II: decreased DTI-p, increased DTI-q (p↓/q); Voxel Group III: increased DTI-p, increased DTI-q (p/q); Voxel Group IV: increased DTI-p, decreased DTI-q (p/q↓). |  |  |  |

In accordance with the univariate histogram features, joint histogram analysis showed that Voxel Group IV (increased DTI-p/decreased DTI-q, p↑/q↓) accounted for the largest proportion in both contrast-enhancing and non-enhancing regions. A significantly higher proportion of Voxel Group IV (p↑/q↓) was found in contrast-enhancing region than non-enhancing region (*p* < 0.001). In contrast, non-enhancing region displayed significantly higher proportion of Voxel Group II (p↓/q↑) and Voxel Group III (p↑/q↑) (both *p* < 0.001) than contrast-enhancing region.

### Multivariate survival analysis

The multivariate survival model of PFS and OS were fitted in the 78 patients for whom all relevant covariates, including IDH-1 mutation, MGMT methylation status were available. The results of the multivariate Cox-regression analysis w ere shown in Table 3.

**Table 3.** Cox multivariate modelling of survivals

| **Table 3. Cox multivariate modelling of survivals** |  |  |  |  |  |  |
| --- | --- | --- | --- | --- | --- | --- |
| Variable | Progression-free survival* |  |  | Overall survival* |  |  |
|  | HR | 95%CI | p value | HR | 95%CI | p value |
| contrast-enhancing region |  |  |  |  |  |  |
| Voxel Group I | 1.02 | 0.99-1.05 | 0.205 | 1.03 | 1.00-1.06 | **0.049** |
| Voxel Group II | 1.06 | 1.004-1.13 | **0.036** | 1.09 | 1.03-1.16 | **0.004** |
| Voxel Group III | 1.01 | 0.99-1.02 | 0.449 | 1.01 | 0.99-1.03 | 0.432 |
| Voxel Group IV | 0.98 | 0.97-1.001 | 0.061 | 0.98 | 0.96-0.996 | **0.015** |
| non-enhancing region |  |  |  |  |  |  |
| Voxel Group I | 1.02 | 0.98-1.06 | 0.463 | 0.997 | 0.95-1.05 | 0.904 |
| Voxel Group II | 1.08 | 1.04-1.13 | **<0.001** | 1.11 | 1.06-1.17 | **<0.001** |
| Voxel Group III | 1.01 | 0.996-1.03 | 0.145 | 1.01 | 0.997-1.03 | 0.116 |
| Voxel Group IV | 0.98 | 0.97-0.997 | **0.014** | 0.98 | 0.97-0.997 | **0.015** |
| *Cox models accounted for sex, age, extent of resection, IDH-1 mutation status, MGMT methylation status and contrast-enhancing tumor volume. HR: hazard ratio; CI: confidence interval; DTI: diffusion tensor imaging; Voxel Group I: decreased DTI-p, decreased DTI-q ($p\downarrow/q\downarrow$); Voxel Group II: decreased DTI-p, increased DTI-q ($p\downarrow/q\uparrow$); Voxel Group III: increased DTI-p, increased DTI-q ($p\uparrow/q\uparrow$); Voxel Group IV: increased DTI-p, decreased DTI-q ($p\uparrow/q\downarrow$). | | | | | | |

Five joint histogram features significantly contributed to both overall survival (OS) and progression-free survival (PFS). Specifically, higher proportion of Voxel Group IV (p↑/q↓) in contrast-enhancing region contributed to a better OS (hazard ratio [HR] = 0.98, *p* = 0.015). Similar contribution to PFS and OS was observed from Voxel Group IV (p↑/q↓) in non-enhancing region (PFS: HR = 0.98, *p* = 0.014; OS: HR = 0.98, p = 0.015). In contrast, higher proportion of Voxel Group II (p↓/q↑) contributed to worse survival: Voxel Group II (p↓/q↑) in the contrast-enhancing region (PFS: HR = 1.06, *p* = 0.036; OS: HR = 1.09, *p* = 0.004); and Voxel Group II (p↓/q↑) in the non-enhancing region (PFS: HR = 1.08, *p* < 0.001; OS: HR = 1.11, *p* < 0.001). The survival curves using Kaplan-Meier method were demonstrated in Figure 2, with *p* values from Log-rank test.

**Figure 2.**
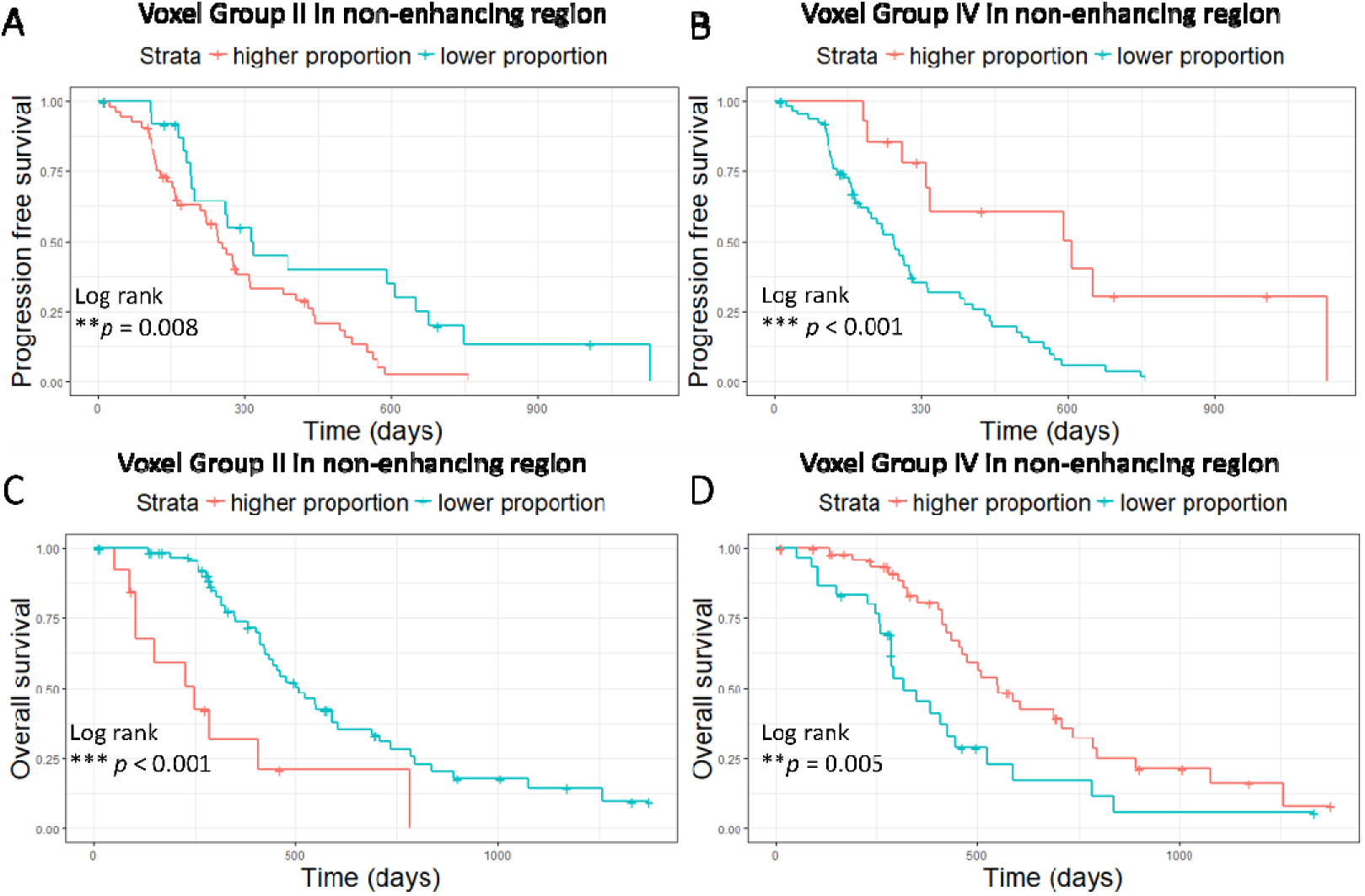
Kaplan-Meier plots of survival analysis. Log-rank test showed lower proportion of Voxel Group II in non-enhancing tumor region was associated with better PFS (p = 0.008) (A) and better OS (p < 0.001) (C). Higher proportion of Voxel Group IV in non-enhancing tumor region was associated with better PFS (p < 0.001) (B) and better OS (p = 0.005) (D).

### Incremental prognostic value of joint histogram features

The results of model comparisons were shown in Table 4. For the prediction of 12-month PFS and OS, the AUC of the baseline multivariate models (including IDH-1 mutation, MGMT methylation status and other clinical factors) were 0.77 (confidence interval [CI]: 0.65-0.88) and 0.81 (CI: 0.70-0.93) respectively. For the prediction of 18-month PFS and OS, the AUC of the baseline multivariate models were 0.75 (CI: 0.61-0.88) and 0.71 (CI: 0.58-0.83) respectively. The incremental prognostic value of joint histogram features was assessed by adding them into the baseline models respectively. Six joint histogram features significantly improved the model (each *p* < 0.05): Voxel group I (p↓/q↓), Voxel group II (p↓/q↑), Voxel group IV (p↑/q↓) in the contrast-enhancing region, and Voxel group II (p↓/q↑), Voxel group III (p↑/q↑), Voxel group IV (p↑/q↓) in non-enhancing region.

**Table 4.** Model Comparison

| **Table 4. Model Comparison** |  |  |  |  |  |  |
| --- | --- | --- | --- | --- | --- | --- |
| Model | 12 month AUC | 95% CI | *p* value | 24 month AUC | 95% CI | *p* value |
| Progression-free survival |  |  |  |  |  |  |
| baseline* | 0.77 | 0.65-0.88 |  | 0.75 | 0.61-0.88 |  |
| Contrast-enhancing region |  |  |  |  |  |  |
| +Voxel Group I | 0.81 | 0.71-0.92 | **0.038** | 0.78 | 0.65-0.92 | 0.424 |
| +Voxel Group II | 0.80 | 0.70-0.91 | 0.153 | 0.86 | 0.76-0.96 | **0.003** |
| +Voxel Group III | 0.77 | 0.66-0.88 | 0.916 | 0.79 | 0.66-0.91 | 0.229 |
| +Voxel Group IV | 0.81 | 0.70-0.91 | 0.077 | 0.81 | 0.68-0.93 | **0.044** |
| Non-enhancing region |  |  |  |  |  |  |
| +Voxel Group I | 0.76 | 0.65-0.88 | 0.792 | 0.76 | 0.63-0.89 | 0.763 |
| +Voxel Group II | 0.80 | 0.70-0.91 | 0.096 | 0.86 | 0.74-0.97 | **0.017** |
| +Voxel Group III | 0.80 | 0.69-0.91 | 0.372 | 0.83 | 0.71-0.95 | **0.023** |
| +Voxel Group IV | 0.80 | 0.70-0.91 | 0.154 | 0.83 | 0.71-0.95 | **0.011** |
| Overall survival |  |  |  |  |  |  |
| baseline* | 0.81 | 0.70-0.93 |  | 0.71 | 0.58-0.83 |  |
| Contrast-enhancing region |  |  |  |  |  |  |
| +Voxel Group I | 0.82 | 0.69-0.95 | 0.070 | 0.75 | 0.63-0.87 | 0.091 |
| +Voxel Group II | 0.84 | 0.73-0.96 | 0.063 | 0.77 | 0.65-0.88 | 0.075 |
| +Voxel Group III | 0.81 | 0.70-0.93 | 0.746 | 0.73 | 0.60-0.85 | 0.443 |
| +Voxel Group IV | 0.84 | 0.72-0.95 | 0.164 | 0.77 | 0.65-0.88 | **0.035** |
| Non-enhancing region |  |  |  |  |  |  |
| +Voxel Group I | 0.81 | 0.70-0.93 | 0.942 | 0.71 | 0.58-0.83 | 0.952 |
| +Voxel Group II | 0.84 | 0.72-0.96 | **0.015** | 0.78 | 0.66-0.89 | **0.022** |
| +Voxel Group III | 0.84 | 0.74-0.94 | 0.109 | 0.74 | 0.61-0.86 | 0.196 |
| +Voxel Group IV | 0.85 | 0.75-0.95 | **0.019** | 0.76 | 0.64-0.88 | 0.053 |
| *Baseline model was built with sex, age, extent of resection, IDH-1 mutation status, MGMT methylation status and contrast-enhancing tumor volume. AUC: area under receiver operator characteristics curve; HR: hazard ratio; CI: confidence interval; DTI: diffusion tensor imaging; Voxel Group I: decreased DTI-p, decreased DTI-q (p↓/q↓); Voxel Group II: decreased DTI-p, increased DTI-q (p↓/q); Voxel Group III: increased DTI-p, increased DTI-q (p/q); Voxel Group IV: increased DTI-p, decreased DTI-q (p/q↓). |  |  |  |  |  |  |

### Stepwise Cox model selection and multivariate prognostic performance

All the clinical factors and the histogram features which significantly improved AUC were tested and selected by the Cox stepwise regression analysis. The results are demonstrated in Table 5. Most significant prognostic variables include extent of resection, MGMT methylation status and Voxel Group II (p↓/q↑) in the non-enhancing region. Specifically, Voxel Group II (p↓/q↑) in the non-enhancing region was significant for both PFS (HR 1.08, *p* < 0.001) and OS (HR 1.36, *p* < 0.001) among all the histogram features.

**Table 5.**
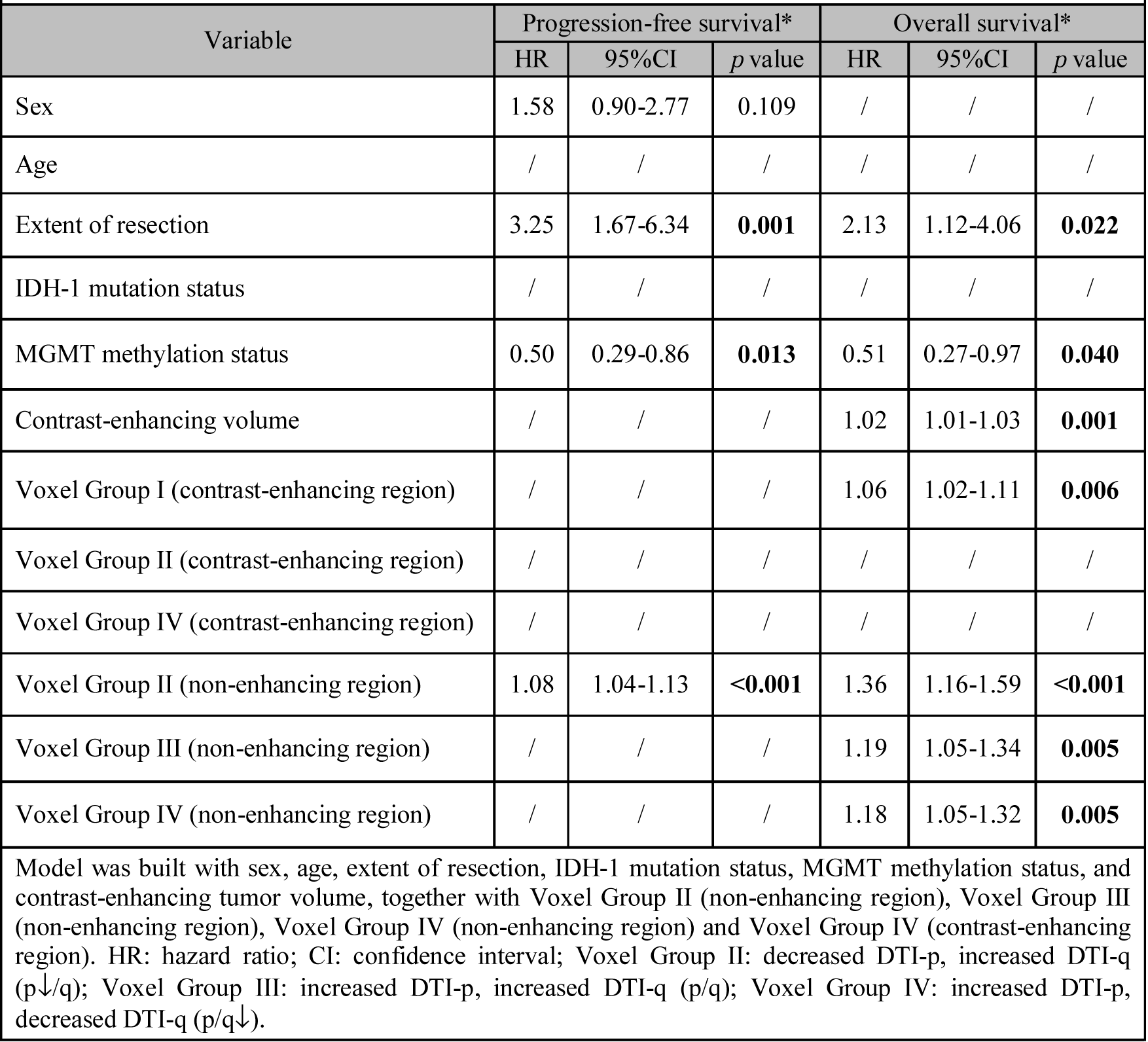
Stepwise Cox multivariate modelling of survivals

| **Table 5. Stepwise Cox multivariate modelling of survivals** |  |  |  |  |  |  |
| --- | --- | --- | --- | --- | --- | --- |
| Variable | Progression-free survival* |  |  | Overall survival* |  |  |
|  | HR | 95% CI | *p* value | HR | 95% CI | *p* value |
| Sex | 1.58 | 0.90-2.77 | 0.109 | / | / | / |
| Age | / | / | / | / | / | / |
| Extent of resection | 3.25 | 1.67-6.34 | **0.001** | 2.13 | 1.12-4.06 | **0.022** |
| IDH-1 mutation status | / | / | / | / | / | / |
| MGMT methylation status | 0.50 | 0.29-0.86 | **0.013** | 0.51 | 0.27-0.97 | **0.040** |
| Contrast-enhancing volume | / | / | / | 1.02 | 1.01-1.03 | **0.001** |
| Voxel Group I (contrast-enhancing region) | / | / | / | 1.06 | 1.02-1.11 | **0.006** |
| Voxel Group II (contrast-enhancing region) | / | / | / | / | / | / |
| Voxel Group IV (contrast-enhancing region) | / | / | / | / | / | / |
| Voxel Group II (non-enhancing region) | 1.08 | 1.04-1.13 | **<0.001** | 1.36 | 1.16-1.59 | **<0.001** |
| Voxel Group III (non-enhancing region) | / | / | / | 1.19 | 1.05-1.34 | **0.005** |
| Voxel Group IV (non-enhancing region) | / | / | / | 1.18 | 1.05-1.32 | **0.005** |
| Model was built with sex, age, extent of resection, IDH-1 mutation status, MGMT methylation status, and contrast-enhancing tumor volume, together with Voxel Group II (non-enhancing region), Voxel Group III (non-enhancing region), Voxel Group IV (non-enhancing region) and Voxel Group IV (contrast-enhancing region). HR: hazard ratio; CI: confidence interval; Voxel Group II: decreased DTI-p, increased DTI-q (p↓/q); Voxel Group III: increased DTI-p, increased DTI-q (p/q); Voxel Group IV: increased DTI-p, decreased DTI-q (p/q↓). |  |  |  |  |  |  |

### Correlations with tumor progression rate and FLAIR volume

Further, we performed correlation tests using above three joint histogram features which showed significantly prognostic values in the stepwise Cox regression analysis. First, the correlation with tumor progression rate was tested in 57 patients had progression and with available MR images at progression. The progression volume (mean ± SD) outside of the resection cavity was 14.3 ± 22.0 cm^3^. The progression rate (mean ± SD) was 0.003 ± 0.013 cm^3^/day. The results showed that Voxel Group II (p↓/q↑) in non-enhancing region had a significant positive correlation (*p* = 0.010, r = 0.35) with the progression rate, whereas Voxel Group IV (p↑/q↓) in non-enhancing region (*p* = 0.040, r = −0.28) showed a negative correlation. No significant correlations were found for Voxel Group III (p↑/q↑) in non-enhancing region.

We analyzed the correlations between the tumor infiltrated volume on FLAIR and the proportions of voxel groups. The results showed that the proportion of Voxel Group II (p↓/q↑) in non-enhancing region showed a significant negative correlation (*p* < 0.001, r = −0.57) with FLAIR volume, whereas the proportion of Voxel Group IV (p↑/q↓) in non-enhancing region (*p* < 0.001, r = 0.37) showed positive correlation. No significant correlation was found with the proportion of Voxel Group III (p↑/q↑) in non-enhancing region. Two examples of pre-operative and progression images, as well as the annotated subregions of Voxel Group II and Voxel Group IV in non-enhancing region are demonstrated in Figures 3 & 4.

**Figure 3.**
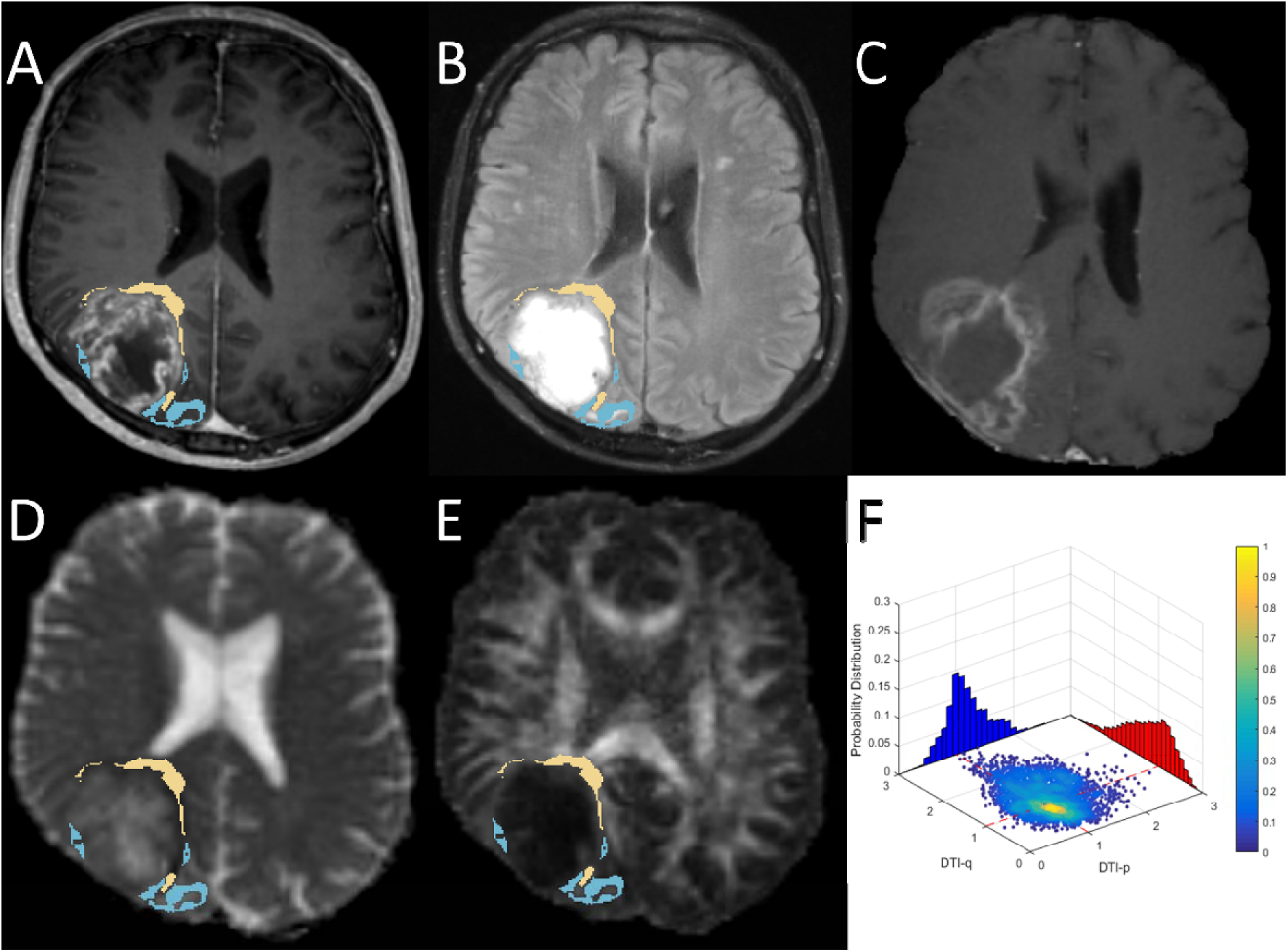
The Voxel Group II (yellow) and Voxel Group IV (blue) of non-enhancing in case 1. The 63-year-old man was radiologically diagnosed with primary glioblastoma (A & B). Volumetric analysis of pre-operative MRI showed contrast enhanced tumor volume was 83.6 cm3. The patient received tumor resection with the guidance of neuro-navigation and 5-aminolevulinic acid fluorescence with the aim of maximal resection, but only subtotal resection was achieved according to 72h post-operative MRI. Pathological assessment confirmed this was a MGMT-methylated glioblastoma and IDH mutation was negative. The patient received concomitant and adjuvant temozolomide chemoradiotherapy. The progression-free survival was 47 days and overall survival was 104 days. The post-contrast T1-weighted imaging showed the progression was around the resection cavity (C). Joint histogram analysis of pre-operative DTI-p (D) and DTI-q (E) maps showed Voxel Group II occupied 15.5% in the non-enhancing tumor and Voxel Group IV occupied 28.2% of the non-enhancing tumor (F).

**Figure 4.**
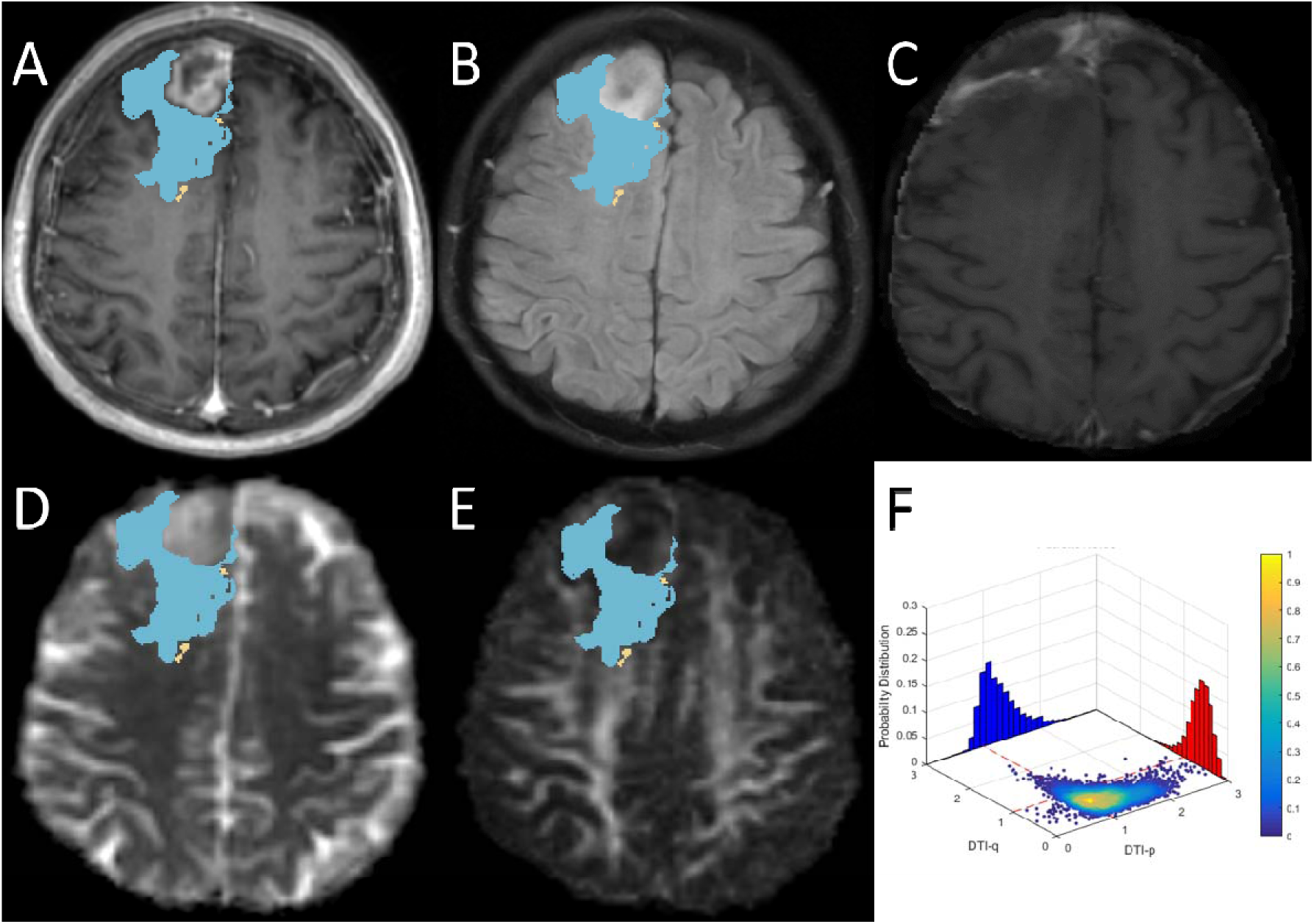
The Voxel Group II (yellow) and Voxel Group IV (blue) of non-enhancing in case 2. The 65-year-old man was radiologically diagnosed with primary glioblastoma (A & B). Volumetric analysis showed contrast enhanced tumor volume was 37.4 cm3. Gross total resection was achieved in this patient with the guidance of neuro-navigation and 5-aminolevulinic acid fluorescence. Pathological assessment confirmed a MGMT-methylated glioblastoma and IDH mutation was negative. The patient received concomitant and adjuvant temozolomide chemoradiotherapy. The progression-free survival was 1006 days and patient was alive in the last follow-up. Post-contrast T1-weighted imaging showed a minor progression around the resection cavity (C). Joint histogram analysis of pre-operative DTI-p (D) and DTI-q (E) maps showed Voxel Group II occupied 2.3% of the non-enhancing tumor and Voxel Group IV occupied 81.5% of the non-enhancing tumor (F).

## Discussion

In this study, we found that preoperative joint histogram analysis using DTI-p and-q can reflect the heterogeneity of tumor infiltration and microstructure change. The histogram features obtained with this method can improve the prognostic value of IDH-1 mutation and MGMT promoter methylation status, and be useful in detection of tumor subregion which might be responsible for progression.

Previous studies have shown that DTI has potential in studying white matter pathology (Zhang, 2010) and is a useful tool to detect tumor infiltration, which causes tissue signature changes (Price et al., 2004; Sternberg et al., 2014). Using stereotactic biopsies, DTI-p and-q is demonstrated to distinguish gross tumor and peritumoral region (Price et al., 2006). As the only in vivo method of describing brain microstructure, it confer additional information for surgical stratification (Jones et al., 2015; Potgieser et al., 2014). However, the interpretation of tensor data is challenging, due to its high dimensionality (Sternberg et al., 2014). Substantial efforts have been made to simplify the tensor data into scalar measures. Among these markers, fractional anisotropy (FA) and mean diffusivity (MD) are most commonly used (Pena et al., 2006). However, since FA can be affected by both the anisotropic and isotropic components (according to its definition equation), its utility is not consistent in detecting tumor infiltration. An enhanced visualization and quantification of tensor imaging was subsequently advanced by decomposing the raw tensor into isotropic (p) and anisotropic (q) components (Wen et al., 2010). This technique has shown its utility in detecting the subtle structure change caused by tumor invasion and predicting progression (Price et al., 2017). Consistent with previous studies, our results showed an increased DTI-p and decreased DTI-q in most univariate histogram features, which have been previously shown to be biomarkers of bulk tumour and invasive tumour components. However, our current results also showed that non-enhancing tumor region may display increased mean and 75th percentile of DTI-q.

One single marker is insufficient to reflect the full tensor. Moreover, it is suggested to combine DTI measures with structural imaging modalities (Bammer, 2003). To achieve this, we proposed the current joint histogram analysis of DTI-p and-q within the tumor regions identified on anatomical sequences. The results showed the Voxel Group IV (p↑/q↓) had the largest proportion. In this subregion, the brain microstructure was displaced to allow more isotropic movement of water molecules. The displacement of fibers also means the ‘fast track’ to infiltrate was diminishing and thus lead to the decreased anisotropic movement. The significantly higher proportion of this diffusion pattern in the bulk tumor than the infiltrated tumor may suggest more substantial fiber damage within the bulk tumor.

More diffusion patterns can be revealed by this approach. Particularly, the higher proportions of Voxel Group II (p↓/q↑) and Voxel Group III (p↑/q↑) in the non-enhancing region may suggest a less disrupted white matter tract. In these regions, water molecules tend to behave higher anisotropic movement. Since the decreased DTI-p is thought to reflect the elevated cell density, the tumor subregion identified by Voxel Group II (p↓/q↑) in non-enhancing region, may represent a migratory tumor habitat with higher cellularity. Though the proportion is relatively low, the contribution to patient survival and tumor progression rate indicated its higher invasiveness and should be treated with more attention in surgical planning. As shown in the case example, some locations of this subregion is in the vicinity of surgical cavity, an extended resection or boosted radiotherapy might be considered for these subregions. Histological correlation of these findings is required.

The joint histogram features found in our study showed clinical significance, with incremental prognostic values when integrated over clinical factors. As the more invasive subregion identified, targeted tumor resection and radiation therapy can perhaps be achieved, which may reduce the radiation damage to the normal brain and enhance the efficacy of treatment. Also, this proposed approach may be applied to a broader imaging research field to meet the demand of multiple modality imaging analysis.

There are some limitations in our study. Firstly, the patient population reported is from a single center and the results were not validated by another cohort. Secondly, although our current study did not have biological validation, previous studies have validated the histological correlates of DTI-p and DTI-q by image-guided biopsies (Price et al., 2006). This current study presented more comprehensive measure of tumor infiltration with the joint analysis of these two directional components.

## Conclusion

We used a novel approach of joint histogram analysis of DTI-p and-q. The results showed that the joint histogram analysis may help to better understand the heterogeneity of tumor infiltration; the decreased DTI-p and increased DTI-q in non-enhancing region may be able to define an invasive subregion responsible for tumor progression. This finding may be useful in targeted surgery and radiation therapy.

## Acknowledgement

We would like to acknowledge the support of National Institute for Health Research, the University of Cambridge, Cancer Research UK and Hutchison Whampoa Limited. The views expressed are those of the author(s) and not necessarily those of the NHS, the NIHR or the Department of Health.

